# SOMSC: Self-Organization-Map for High-Dimensional Single-Cell Data of Cellular States and Their Transitions

**DOI:** 10.1101/124693

**Authors:** Tao Peng, Qing Nie

**Affiliations:** Department of Mathematics, University of California Irvine, Irvine, 92697, USA; Department of Developmental and Cell Biology and Department of Biomedical Engineering, University of California Irvine, Irvine, 92697, USA

## Abstract

Measurement of gene expression levels for multiple genes in single cells provides a powerful approach to study heterogeneity of cell populations and cellular plasticity. While the expression levels of multiple genes in each cell are available in such data, the potential connections among the cells (e.g. the cellular state transition relationship) are not directly evident from the measurement. Classifying the cellular states, identifying their transitions among those states, and extracting the pseudotime ordering of cells are challenging due to the noise in the data and the high-dimensionality in the number of genes in the data. In this paper we adapt the classical self-organizing-map (SOM) approach for single-cell gene expression data (SOMSC), such as those based on single cell qPCR and single cell RNA-seq. In SOMSC, a cellular state map (CSM) is derived and employed to identify cellular states inherited in the population of the measured single cells. Cells located in the same basin of the CSM are considered as in one cellular state while barriers among the basins in CSM provide information on transitions among the cellular states. A cellular state transitions path (e.g. differentiation) and a temporal ordering of the measured single cells are consequently obtained. In addition, SOMSC could estimate the cellular state replication probability and transition probabilities. Applied to a set of synthetic data, one single-cell qPCR data set on mouse early embryonic development and two single-cell RNA-seq data sets, SOMSC shows effectiveness in capturing cellular states and their transitions presented in the high-dimensional single-cell data. This approach will have broader applications to analyzing cellular fate specification and cell lineages using single cell gene expression data

## 1. Introduction

Heterogeneity of cell populations is considered functionally and clinically significant in normal and diseased tissues, and transitions among different subpopulations of cells play key roles in cell differentiation during development or disease recurrence (Tsioris *et al.*, 2014; Wilson *et al.*, 2014; Saadatpour *et al.*, 2015). In recent years, single-cell gene expression profiling technologies have emerged as an important tool in dissecting heterogeneity and plasticity of cell populations and in analysis of cell-to-cell variability on a genomic scale (Saliba *et al.*, 2014). For example, mammalian pre-implantation development has been analyzed from oocyte stage to morula stage in both human and mouse embryos using single-cell RNA sequencing (Xue *et al.*, 2013; Yan *et al.*, 2013) to identify stage-specific transcriptomic dynamics; In breast cancer, gene expression profiles of tumor subpopulations along a spectrum from low metastatic burden to high metastatic burden have been obtained using qPCR at the single-cell level (Lawson *et al.*, 2015); and multiple new phenotypes in healthy and leukemic blood cells have been identified using gene expression signatures through analysis of single-cell data (Levine *et al.*, 2015).

Distinguishing or clustering measured cells computationally through their transcriptomic data (e.g. gene expression) is challenging. The number of genes measured is usually significantly larger than the number of cells (Jiang *et al.*, 2004). Another challenge is that a group of cells collected at one temporal point from one sample may not be perfectly ordered in time compared to the cells collected at a slightly different temporal stage, due to cell-to-cell variability in sampling and the nature of unsynchronized cell divisions (Heppner, 1984; de Vargas Roditi and Claassen, 2015). As a result, the pseudotemporal ordering of single cells in a high-dimensional gene expression space was introduced (Trapnell *et al.*, 2014b). The difficulty in analyzing singlecell data becomes particularly evident for systems of differentiation in which new cell types emerge as time advances, such as identifying lineage specific markers of different cell subtypes during the development of murine lung (Treutlein *et al.*, 2014) and finding the differentiation trajectory of skeletal muscles (Trapnell *et al.*, 2014a).

Temporal ordering of single cells, grouping cells of similar transcriptomic profiles, finding transition points, and determining branches are the key steps in analyzing the single-cell data. Clustering methods based on Principle Component Analysis (PCA) or Independent Components Analysis (ICA), such as MONOCLE algorithm (Trapnell *et al.*, 2014a), allow grouping cells according to the specific properties of interest. Several other clustering-based methods such as SPADE (Qiu *et al.*, 2011), t-SNE (Van der Maaten and Hinton, 2008), and viSNE (Amir *et al.*, 2013) were introduced to identify subpopulations within the measured cells without an explicit temporal ordering of cells. In the Wanderlust algorithm (Bendall *et al.*, 2014), a pseudo-temporal ordering technique incorporated the continuity concept in branching process, however, with an assumption that the cells consist of only one branch during differentiation. To study nonlinearity of the branching process in differentiation, a diffusion map technique was adapted to single-cell data by adjusting kernel width and inclusion of uncertainties, enabling a pseudo-temporal ordering of single cells in a high-dimensional gene expression space (Haghverdi *et al.*, 2015). SLICER is another method to capture highly nonlinear gene expression changes and select genes related to the process, and to detect multiple branches (Welch *et al.*, 2016). TASIC was developed to determine temporal trajectories, branching and cell assignments using a probabilistic graphical model (Rashid *et al.*, 2017). UNCURL incorporates prior knowledge to perform the cell state identification (Mukherjee *et al.*, 2017). With a focus on modeling the dynamic changes associated with cell differentiation, a bifurcation analysis method (SCUBA) was developed to extract lineage relationships (Marco *et al.*, 2014).

The Waddington landscape (Goldberg *et al.*, 2007) of gene expression provided a global and convenient view in describing stem cell dynamics and lineages. In this method, a forward stochastic model on a small gene network was first derived, then a landscape of cellular states was obtained by constructing an energy function that depends on each gene in the modeled regulatory network (Foster *et al.*, 2009; Zhang *et al.*, 2013; Zhou and Huang, 2011; Chen *et al.*, 2015). The prior knowledge of the gene regulatory network needs to be known in this method, and the landscape calculation did not require dimension reduction in the gene space. However, due to the computational cost associated with sampling solutions of stochastic differential equations or solving equations of probability density functions of the gene states, the network size in the landscape calculation can’t be too large (Wang *et al.*, 2011).

Here, we propose a new method to analyze single-cell gene expression data by combining a machine learning method and a concept similar to the landscape (Wang *et al.*, 2011; Kohonen, 1998) (Figure 1). In this approach, the high-dimensional data of single cells is first reduced to two dimensions through a classical unsupervised artificial neural network (ANN) method: a self-organizing map (SOM) (Kohonen, 1998) in which the topological properties of the input data are preserved through a neighborhood function. The cellular states are then identified by the watershed algorithm based on the Umatrix calculated by SOM (Vincent and Soille, 1991; Najman and Schmitt, 1996). By building transition paths among the cellular states, we obtain a cellular state map (CSM). In this map, the barriers separating different states provide information on transitions between cellular states. Moreover, a replication probability and transition probabilities are estimated in the cellular state transition paths. Next, the state-driven genes differentially expressed during a cellular state transition are determined by t-test, and then pathway enrichment analysis for each transition is performed based on the list of state-driven genes. In this approach, the transition path among the cellular states leads to a pseudo-temporal ordering of the cells. To study effectiveness and capability of the approach, we apply SOMSC to a set of simulated data and three real data sets based on qPCR or RNA-seq collected from cell at various stages of differentiation.

**Figure 1.**
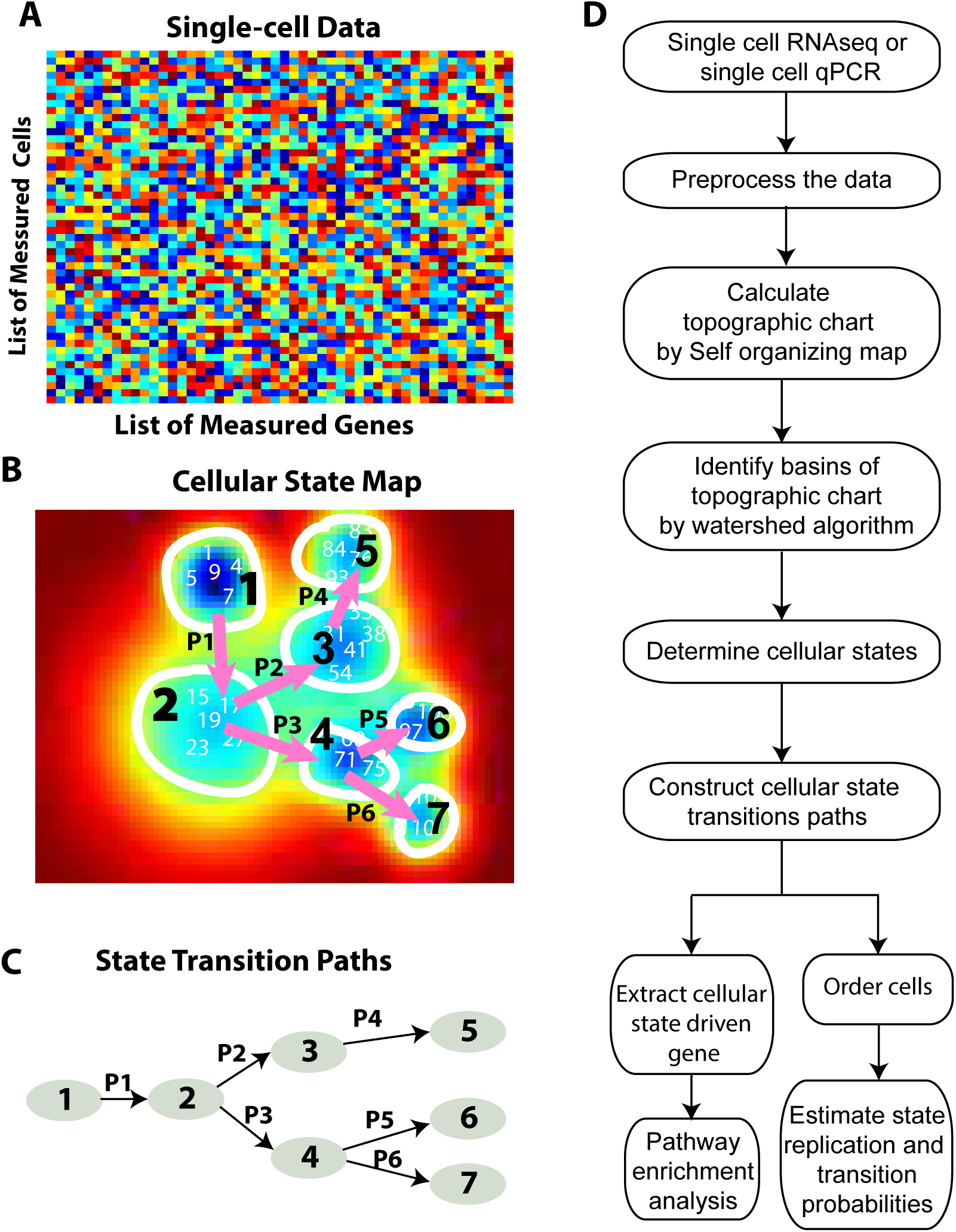
A schematic diagram on steps of constructing cellular state maps and transition paths using the SOMSC method. (A) The gene expression data of single cells. (B) A topographic chart constructed by SOMSC using the data. The transitions among different basins are labeled by arrows: P1, P2, …, and P5. (C) The cellular state lineage trees or differentiation processes are then summarized based on the transition paths. (D) The workflow of SOMSC.

## 2. Methods

SOMSC has three major functions: identifying cellular states, reconstructing cellular state transition paths and building the pesudotime ordering of cells. The algorithm of SOMSC consists of six main steps as follows (Figure 1).

### Step 1: Calculate a topographic chart of single cell data using a self-organizing map

SOMSC takes single-cell RNAseq and single-cell PCR data as the inputs *G* = (*g*_1_*, g*_2_*, …, g*_*N*_)^*T*^, where *g*_*i*_ is the vector with the length *n* of the gene expression levels for the *i*-th sample and *N* is the number of the samples. Since two kinds of datasets are collected using different technology platforms, different preprocessing methods are necessary (See details in Supplementary file). A topographical chart of high dimensional expression data is calculated by a SOM. A SOM is an effective way of analyzing topology of high-dimensional data by projecting the data to low-dimensional surfaces through a rectangular, a cylindrical, or a toroidal map (Kohonen, 1998). In the SOM, an ordered set of model vectors *x ∈ R*^*n*^ is mapped onto the space of observation vectors *m*_*i*_*∈ R*^*n*^ through the following iteration processes:

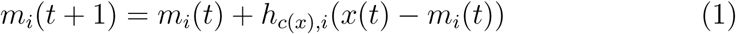

where *t* is a regression index. This regression procedure is performed recursively for each sample *x*(*t*). The scalar multiplier *h*_*c*(*x*)*,i*_ is a Gaussian neighborhood function, acting like a smoothing or blurring kernel over SOM computational grids:

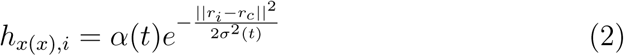

where 0 *< α*(*t*) *<* 1 is a learning-rate factor, which decreases monotonically in each regression step; *r*_*i*_*∈R*^2^ and *r*_*c*_*∈ R*^2^ are the computational grid locations, and *σ*(*t*) corresponds to the width of the neighborhood function that also decreases monotonically in each regression step. The subscript *c* = *c*(*x*) is obtained when the following condition is achieved:

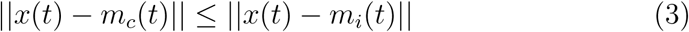

Consequently, *m*_*c*_(*t*) is the “winner” that matches the best with *x*(*t*). The comparison metric *|| • ||* is selected as the Euclidean metric for this study in Eq. 2, and Eq. 3. If multiple functions *c*(*t*) satisfy Eq. 3 with discrete-valued variables, *c*(*t*) is chosen randomly among those functions for the winner. In addition, a toroid map is used to reduce edge effects in the data on the overall mapping (Vesanto *et al.*, 1999). Applying the SOM to the single-cell gene expression data leads to a unified distance matrix (U-matrix) *U* (*x, y*), representing distances between neighboring map units, where *x* and *y* denote the coordinates of a plane (Kohonen, 1998). Accordingly, *U* (*x, y*) defines a 2-dimensional topographic chart.

### Step 2: Identify basins of the topographic chart by the watershed algorithm

The watershed segmentation algorithm is employed to identify the basins of the topographic chart. By the SOM the high dimensional single cell data is projected onto *U* (*x, y*) denoting the distance between adjacent grids. The algorithm identifies the boundaries of basins by constantly pouring water into the mountains of the chart (Najman and Schmitt, 1994). As water levels rise, the boundaries of basins are estimated (See more details of the watershed segmentation algorithm in Supplementary file). Let *nb* denote the number of basins in the topographic chart, and *C*_1_, *C*_2_, *…*, *C*_*nb*_ represent the identified basins. The height of the barrier between two adjacent basins is the number of genes differentially expressed in those cells of the two adjacent basins, which are calculated by t-test with 5% significance. The adjacent matrix of basins is then calculated: *T* = (*t*_*ij*_)_*n,n*_, where *t*_*ij*_ is the height of the barrier between *C*_*i*_ and *C*_*j*_ and *t*_*ij*_ = 0 means that *C*_*i*_ and *C*_*j*_ are not adjacent with each other.

### Step 3: Classify the cellular states and construct their transition paths

The cellular states are classified based on the adjacent matrix calculated in Step 2. In order to avoid the over-segmentation occurrence when applying the watershed algorithm, we merge any two basins *C*_*i*_ and *C*_*j*_ when *C*_*i*_ and *C*_*j*_ form a cycle, which means the height of the barrier between these two basins is the smallest one between the basin *C*_*i*_ and all its adjacent basins as well as the smallest one between the basin *C*_*j*_ and all its adjacent basins. (*C*_*i*_,*C*_*j*_) is called a merging pair. After the merging, all cells in one basin labeled as *S*_*i*_, (*i* = 1,*…*,*m*) are in the same cellular state. The adjacency matrix *D* =(*d*_*ij*_)_*m,m*_ represents the distances between the cells in *S*_*i*_ and the ones in *S*_*j*_ (the default distance metric in our algorithm is the *l*_1_ norm), and can be considered as an undirected network for cellular states in SOMSC, which is the cellular state network. All nodes in the network are labeled by the cellular states *S*_*i*_ (*i* = 1,2,*…*,*m*). The weight of the edge connecting the node *S*_*i*_ and the node *S*_*j*_ is *d*_*ij*_ in the adjacency matrix *D* and there is no edge between the node *S*_*i*_ and the node *S*_*j*_ if *d*_*ij*_ = 0. All edges in the cellular state network are denoted by *F*. We assume that the starting cellular state is *S*_*s*_ where *s ∈* {1, 2, *…*, *m*}. First, we find out the edge pair in the path (*S*_*i*_, *S*_*i*_*p*__) for each node *S*_*i*_ where *i ∈* {1, 2, *…*, *m*}. *S*_*i*_*p*__ is the closet adjacent cellular state of *S*_*i*_. All edge pairs are denoted by *E*. Second, we make all nodes of the cellular states reachable. A node *S*_*i*_ is reachable when there exists a path from *S*_*i*_ to the node *S*_*s*_ of the starting cellular state, which consists of the edge pairs in *E*. Here we denote the set of all reachable nodes by *R* and the set of the rest nodes by *Q*. If the set *Q* is not empty, then we will find out the edges from *F\E* (the element in *F* but not in *E*) which connect the node in *R* and the one in *Q* and the one with smallest weight is selected to add to the set *E* and the sets of *R* and *Q* are updated correspondingly. We repeat these steps until *Q* is empty since each iteration results in more reachable nodes in *R*. Accordingly, all nodes become reachable with the edge pair set *E*, and the path *P*_*i*_ from *S*_*i*_ to *S*_*s*_ consists of a subset of the edge pair set *E*. Finally, all the paths *P*_*i*_ (*i* = 1,2,*…*,*m*) are combined to generate a transition path tree of the cellular states. From 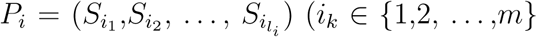 we can extract the transition pairs 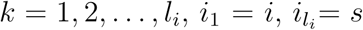, which mean that the cellular state 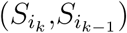 is the parent of *S*_*i*_*k-*1__. All transition pairs determine the transition path tree (*S*_*p*_1__, *…*, *S*_*p*_*m*__) together, where *S*_*p*_*i*__ is the parent cellular state of *S*_*i*_ in the path tree and *S*_*i*_ is the daughter cellular state of *S*_*p*_*i*__.

### Step 4: Construct the cellular state map for all cells

Each cell in the data is assigned a relative location in the cellular state map using the following method. Not all cells in the data show up in the topographic chart since only winners of the grids in the SOM stay, suggesting known cellular states of those cells. We then use k-nearest neighbors (KNN) algorithm to identify the states of the remaining cells (Altman, 1992).

To visualize the topologic structure of the data we define the cellular state map (CSM) as follows. First, we calculate the convolution function *M* by the following equations.

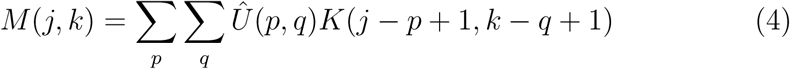

where

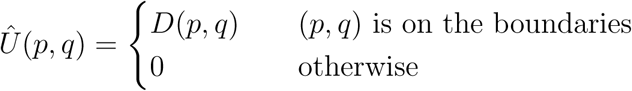

*K* is a kernel matrix whose elements define how to remove the high frequency components of the original data *Û*. The size of kernels may be different from the size of the *Û*. Small-sized kernels can smooth data containing only a few frequency components whereas larger size kernels can provide better precision for tuning frequency response, resulting in a smoother output. Accordingly, the CSM is the central part of the convolution *M* whose size is equal to the size of *Û*. Each cell is plotted as a three-dimensional sphere, whose center represents that cell’s position in the CSM.

### Step 5: Detect the state-driven genes and their enrichment analysis in each transition

We next identify the differentially expressed genes for each cellular state transition. We perform a t-test for the expression levels of each gene in all cells involved in the cellular state transition. A gene is taken to be state-driven if the p-value is within 1%. Accordingly, a list of all state-driven genes for each cellular state transition can be obtained. The gene list is then used as an input into the Enrichr database (http://amp.pharm.mssm.edu/Enrichr/), which defines the members of each pathway as a gene list. Fisher’s exact test is used to obtain a p-value for the number of pathway members present among the state-driven genes we obtained. If the p-value falls within our critical region (5%), we determine the pathway to be significantly enriched (Chen *et al.*, 2013).

### Step 6: Estimate the cellular state replication probability and cellular state transition probabilities and determine the pseudo-time ordering of each cell during the cellular state transitions

One goal of SOMSC is to estimate the cellular state replication probability and the cellular state transition probability for each cellular state. First, we calculate the centroid for each cellular state and denote *CT*_*i*_ as the centroid of the cellular state *S*_*i*_. Second, we define the cellular state replication axis *P R*_*i*_ and the transition axes 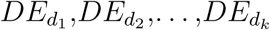 for the corresponding *k* daughter cellular states 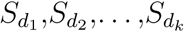. They are calculated by *P R*_*i*_ = *CT*_*p*_–*CT*_*i*_ and 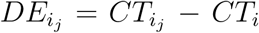 where *j* = 1,2,*…*,*k*; *CT*_*p*_ is the centroid of the parent cellular state of *S*_*i*_; and 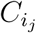 is the centroid of the *j*th daughter cellular state of *S*_*i*_. Then we project the data 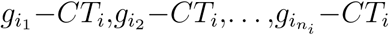, where *n*_*i*_ is the number of cells in the cellular state *S*_*i*_, to the cellular state replication axis and the cellular state transition axises. Third, we obtain the vector 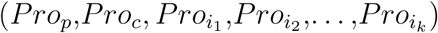 for the expression vector of each cell 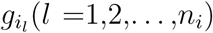, where *P_ro_p__* is the projection of the data of the *l*-th cell in the cellular state *S*_*i*_ to the cellular state replication axis *PR*_*i*_, *P_ro_c__* is the distance between 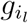 and *CT*_*i*_, and 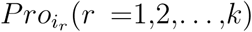 is the projection of *g*_*i*_*l*__ to the cellular state transition axis 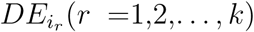. We then determine the replication/transition state of the cells as follows: If the minimum positive element in 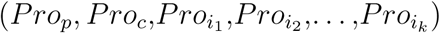 is the projection to one cellular state transition axis 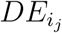, then it means the *l*-th cell will transition to the daughter cellular state 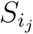. Otherwise, the *l*-th cell is in the replication cellular state. The number of cells in the replication and transition states are obtained and the normalized numbers are taken as the replication and transition probabilities, respectively. Fourth, the pseudotime ordering of each cell is determined by its projection to the replication axis or the transition axes or the distance from the cell to the centroid of the cellular state *S*_*i*_. If the cell *g*_*i*_*l*__ is in the replication state and *P_ro_p__* is positive, then we get the ordering pair (1, – *P_ro_p__*). If the cell *g*_*i*_*l*__ is in the replication state and *P_ro_p__* is negative, then we get the ordering pair (2, *P_ro_c__*). If the cell *g*_*i*_*l*__ is in the transition state, then we get the ordering pair (3, *P*_*ro*_min__), where *P*_*ro*_min__ is the minimum positive element of 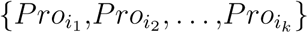. In this way, we get the ordering pair for each cell in the cellular state *S*_*i*_. The pseudotime ordering of cells in *S*_*i*_ is based on the ordering pairs. We compute the pseudotime ordering of cells in different cellular states based on the cellular transition path. Finally, we construct the trajectories of the expression levels of each gene from all cells along the pseudotime ordering. Here we interpolate the average expression levels of the gene in disjoint small groups of cells along the pseudotime ordering.

#### Generate the simulation data

In order to effectively evaluate performance and choices of parameters of SOMSC, we next construct a toy system consisting of a small number of genes to mimic single-cell gene expression data. There are three stages in the system, and in each stage one type of cells makes a transition to two other types of cells (Figure 2A). Together, seven types of cells with three branches are present in the system. The cellular types are defined by the expression levels of six genes (Figure 2A). Specifically, in Type 1 cells Gene A and Gene B are activated and the other four genes are silenced; in Type 2 cells Gene A, Gene C, and Gene D are activated; in Type 3 cells Gene B, Gene E, and Gene F are activated; when one of Gene A and Gene B and one of Gene C, Gene D, Gene E and Gene F are activated, four other types of cells in the third stage are then defined as Type 4, Type 5, Type 6, and Type 7 cells, respectively.

**Figure 2.**
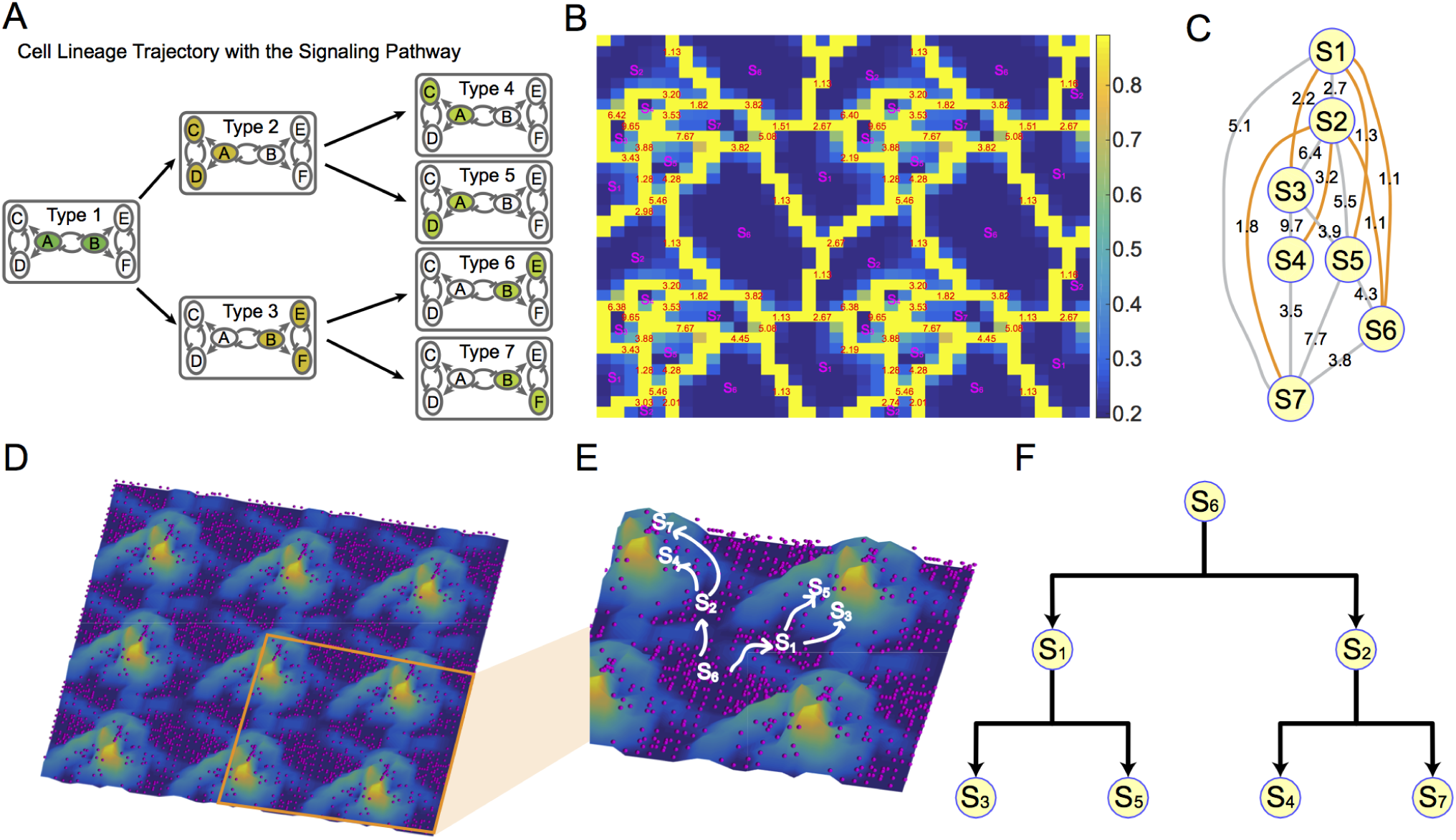
SOMSC on the simulated model. (A) A three-stage lineage system. Stage 1 contains one type of cells in which the activated genes, A and B are highlighted by green; Stage 2 contains Type 2 cells and Type 3 cells in which the activated genes, A, C, and D are highlighted by orange in Type 2 cells while the activated genes, B, E, and F are highlighted by orange in Type 3 cells; Stage 4 contains four types of cells: Type 4 cells, Type 5 cells, Type 6 cells, and Type 7 cells. The activated genes, A and C, A and D, B and E, or B and F are highlighted in light green in Type 4, Type 5, Type 6, and Type 7 cells, respectively. (B) The topographic chart is constructed based on SOMSC algorithm with the map of 18*×*18 grids in the topographic chart computed for *N* = 346 single cells. The basins are labeled by *S*_1_, *S*_2_, *S*_3_, *S*_4_, *S*_5_, *S*_6_, *S*_7_. Each basin represents a cluster of cells in one cellular state. The yellow areas are the boundaries between adjacent basins and the red numbers are the heights of the barriers of adjacent basins. (C) The cellular state network is built based on the topographic chart. The nodes are the cellular states. The weight of the edge is the height of the barrier of adjacent basins. The orange line is the edge with the smallest weight associated with each cellular state. (D) The cellular state map. The red dots are the cells. The basins correspond to the cellular states in Figure B. (E) The zoomed-in cellular state map. The white text is the label of the cellular state. The arrow is the direction of the cellular state transition. (F) The cellular state transition map. The percentage numbers present the probability of the cellular state transition replication and cellular state transitions.

The system of three-toggle modules consisting of six genes is modeled through a system of stochastic differential equations (Haghverdi *et al.*, 2015; Chen *et al.*, 2005; Ocone *et al.*, 2015). Starting with only Type 1 cells in the system (i.e. the initial state), the expression values of each gene are then collected at three different temporal stages for each stochastic simulation: the early, the middle, and the final stage, in order to mimic a typical set of temporal single-cell data (See Section II in the Supplementary file). Repeating the stochastic simulations using the same set of parameters and the same initial values of genes for 400 times produces a set of gene expression values, corresponding to 1200 sets of single-cell data.

## 3. RESULTS

### 3.1. SOMSC on the simulation data

We apply SOMSC to 346 cells, which are randomly selected from 1200 simulated cells. The SOM (*N* = 16) maps the high-dimensional geneexpression data to a 2-dimensional topographic chart, *U* (*x, y*), and the watershed algorithm is employed to identify the basins of the topographic chart with the yellow boundaries between the adjacent basins (Figure 2B). There are seven basins identified. Since no cycles exist on the topographic chart, basins correspond to cellular states, which are labeled *S*_1_, *S*_2_,*…*,*S*_7_ (Figure 2B). The red numbers on the yellow boundaries are the height of the boundaries between the adjacent cellular states. Based on the above information, we calculate the adjacency matrix of cellular states, *T* = (*t*_*ij*_)_7,7_ and construct a cellular state network, in which the edge between the node *S*_*i*_ and the node *S*_*j*_ means that *t*_*ij*_*≠* 0 and the weight of the edge is *t*_*ij*_ (Figure 2C). Then we identify the edge pair for each cellular state (Table 1), highlighted in orange (Figure 2C). For example, the cellular state *S*_1_ has five adjacent cellular states, *S*_2_, *S*_3_, *S*_5_, *S*_6_, and *S*_7_ with edge weights, 2.6733, 2.1888, 1.2798, 1.1335, 5.0827 and *S*_6_ is the one with the smallest weight, 1.1335. Therefore, (*S*_1_, *S*_6_) is the edge pair for the cellular state *S*_1_.

**Table 1:** Edge pairs in the simulation data

| Cellular state | Edge in the path | Cellular state | Edge in the path |
| --- | --- | --- | --- |
| $S_1$ | $(S_1, S_6)$ | $S_2$ | $(S_2, S_6)$ |
| $S_3$ | $(S_3, S_1)$ | $S_4$ | $(S_4, S_2)$ |
| $S_5$ | $(S_5, S_1)$ | $S_6$ | $(S_6, S_1)$ |
| $S_7$ | $(S_7, S_2)$ | | |

We track the paths from each cellular state to the starting cellular state *S*_6_ using the highlighted edges. Since the edge pair for the cellular state *S*_1_ is (*S*_1_, *S*_6_), the transition path *P*_1_ is from the cellular state *S*_1_ to the cellular state *S*_6_, (*S*_1_, *S*_6_). Similarly, the transition path *P*_2_ is from the cellular state *S*_2_ to the cellular state *S*_6_, (*S*_2_, *S*_6_). Because the edge pair for the cellular state *S*_3_ is (*S*_3_, *S*_1_) and the path transition *P*_1_ is (*S*_1_, *S*_6_), then the combination of (*S*_3_, *S*_1_) and *P*_1_ results in the transition path *P*_3_, (*S*_3_, *S*_1_, *S*_6_). Similarly, We find the transition paths *P*_4_, *P*_5_, *P*_7_ from these cellular states *S*_4_, *S*_5_, *S*_7_ to the starting cellular state *S*_6_. Those transition paths are summarized as follows (Table 2).

**Table 2:** Path from one cellular state to the starting cellular state in the simulation data

| State | Path | The path to the starting cellular state |
| --- | --- | --- |
| $S_1$ | $P_1$ | $(S_1, S_6)$ |
| $S_2$ | $P_2$ | $(S_2, S_6)$ |
| $S_3$ | $P_3$ | $(S_3, S_1, S_6)$ |
| $S_4$ | $P_4$ | $(S_4, S_2, S_6)$ |
| $S_5$ | $P_5$ | $(S_5, S_1, S_6)$ |
| $S_6$ | $P_6$ | $(S_6, S_6)$ |
| $S_7$ | $P_7$ | $(S_7, S_2, S_6)$ |

The cellular state map is constructed to illustrate the relative locations of all cells and the relationship among different cellular states (Figure 2D) (See Methods). The SOM maps the winner of each computational grid to the topographical chart and the KNN algorithm places each non-winner cell at the mean position of its nearest neighbors. Cells are represented on the topographic chart as purple spheres (Figure 2D). Finally, we add the arrows from the parent cellular state to the daughter cellular state on the CSM (Figure 2E). The cellular state transition tree is obtained from all transition pairs, (*S*_1_, *S*_3_), (*S*_1_, *S*_4_), (*S*_3_, *S*_5_), (*S*_3_, *S*_7_), (*S*_4_, *S*_2_), and (*S*_4_, *S*_6_) (Figure 2F). To validate the calculated transition path we plot the cellular states of the cells in the topographic chart (Figure S4). All above results show that SOMSC can not only calculate the transition path but also identify the cellular states of over 90% cells.

### 3.2. SOMSC on qPCR data of mouse embryo development from zygote to blastocyst

Previously, the expression levels of 48 genes at seven time points were measured using qPCR for mouse early embryonic development from zygote to blastocyst (Guo *et al.*, 2010). Expression levels for each of the 439 singlecell libraries were normalized independently by the mean expression levels of two genes: Actb and Gapdh (Guo *et al.*, 2010).

Two different approaches might be applied to such data by either using the data at each temporal point individually or lumping the data of all seven stages into one set. However, a series of single-state maps is unable to determine potential cellular state transition paths from the data because different basins or cellular states are obtained using different topographic charts (Figure S5). Therefore we choose to analyze all time points concurrently. Furthermore, prior knowledge about cell state is withheld during inference and subsequently used to validate the state-transition path calculated by SOMSC.

We generate a topographic chart from the expression-levels of all 439 cells and use the watershed algorithm to identify 20 basins. The basins are labeled by *C*_1_, *C*_2_, *…*, *C*_20_ and the heights of the barriers between adjacent basins are the red numbers on the yellow barriers (Figure S6 and S7). The t-test is performed gene-by-gene between the cells of each basin-pair, *C*_*i*_ and *C*_*j*_ and then we calculate the number *T*_*ij*_ of the genes, which are not differentially expressed at 5% significance. Based on all calculated *T*_*ij*_, we merge the following pairs: (*C*_13_, *C*_14_), (*C*_17_, *C*_18_), (*C*_18_, *C*_20_), (*C*_1_, *C*_6_), (*C*_2_, *C*_3_), (*C*_15_, *C*_16_), (*C*_4_, *C*_7_) (See Methods), yielding a new topographic chart with barriers highlighted in yellow (Figure 3B). Thirteen basins labeled by *S*_1_, *S*_2_, *…*, *S*_13_ are identified in the new chart, corresponding to thirteen cellular states. The distance between adjacent states is labeled in red on the yellow boundaries between states (Figure 3B, S8 and S9).

**Figure 3.**
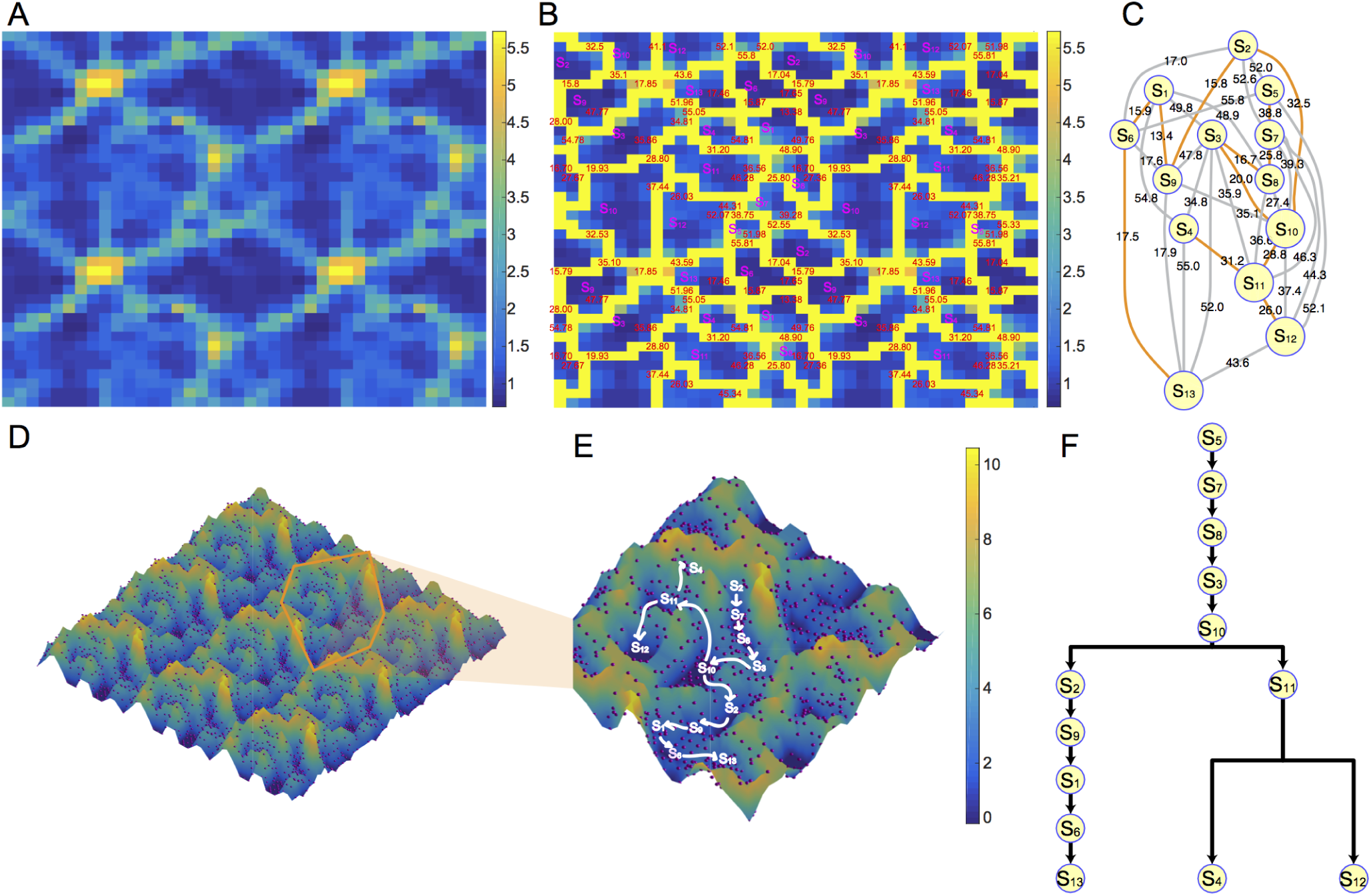
SOMSC reconstructed the cellular state transition path using the qPCR data of mouse stem cells from zygote to blastocyst (Guo *et al.*, 2010). (A) The topographic chart is constructed based on the SOMSC algorithm with the map of 20*×*20 grids. (B) The topographic chart with barriers of the adjacent cellular states. The yellow areas are the barriers of the adjacent cellular states. Each basin means one cellular state: *S*_1_, *S*_2_, *…*, *S*_13_. The red numbers are the heights of the barriers. (C) The cellular state network is built based on the topographic chart. The nodes are the cellular states. The weight of the edge is the height of the barrier of adjacent basins. The orange line is the edge with the smallest weight associated with each cellular state. (D) The cellular state map. The red dots are the cells. The basins correspond to the cellular states in Figure B. (E) The zoomed-in cellular state map. The white text is the label of the cellular state. The arrow is the direction of the cellular state transition. (F) The cellular state transition map. The percentage numbers present the probability of the cellular state transition replication and cellular state transitions.

Using the topographic chart, we construct a cellular state transition path tree in two steps. First, we construct the cellular state network (Figure 3C) and highlight the edges with the smallest weight values for each cellular state. The highlighted edges for the cellular states are summarized in Table 3.

**Table 3:** Edge pairs in the qPCR data of mouse embryo development from zygote to blastocyst

| Cellular state | Edge in the path | Cellular state | Edge in the path |
| --- | --- | --- | --- |
| $S_1$ | $(S_1, S_9)$ | $S_2$ | $(S_2, S_9)$ |
| $S_3$ | $(S_3, S_8)$ | $S_4$ | $(S_4, S_{11})$ |
| $S_5$ | $(S_5, S_7)$ | $S_6$ | $(S_1, S_6)$ |
| $S_7$ | $(S_7, S_8)$ | $S_8$ | $(S_3, S_8)$ |
| $S_9$ | $(S_1, S_9)$ | $S_{10}$ | $(S_3, S_{10})$ |
| $S_{11}$ | $(S_{11}, S_{12})$ | $S_{12}$ | $(S_{11}, S_{12})$ |
| $S_{13}$ | $(S_6, S_{13})$ | | |

Second, we track the cellular state transition paths from each cellular state to the starting cellular state *S*_5_ using the highlighted edges. We find the paths from these cellular states *S*_3_, *S*_4_, *S*_7_, *S*_8_, *S*_10_, *S*_11_, *S*_12_ to the starting cellular state *S*_5_ consisting of the edges from Table 3. They form one group *R* of reachable cellular states and the rest cellular states *S*_1_, *S*_2_, *S*_4_, *S*_6_, *S*_9_, *S*_11_, *S*_12_, *S*_13_ form the group *Q* using the highlighted edges. Then the smallest weight value between *R* and *Q* is the one, 28.797, between the cellular state *S*_11_ and the cellular state *S*_10_. Then *R∪S*_11_*∪S*_4_*∪S*_12_ generates the new *R* and the new *Q* is the set consisting of *S*_1_, *S*_2_, *S*_6_, *S*_9_, and *S*_13_. The edge with the smallest weight, 32.526, connecting *R* and *Q* is (*S*_2_, *S*_10_), which results all elements in *Q* are reachable. Accordingly, all these edges (*S*_1_, *S*_9_), (*S*_2_, *S*_9_), (*S*_3_, *S*_8_), (*S*_4_, *S*_11_), (*S*_5_, *S*_7_), (*S*_1_, *S*_6_), (*S*_7_, *S*_8_), (*S*_3_, *S*_10_), (*S*_11_, *S*_12_), (*S*_6_, *S*_13_), (*S*_2_, *S*_10_), and (*S*_10_, *S*_11_) result in that each cellular state can reach the starting cellular state *S*_5_ by a transition path (Table 4).

**Table 4:** Path from one cellular state to the starting cellular state in the qPCR data of mouse embryo development from zygote to blastocyst

| State | Path | The path to the starting cellular state |
| --- | --- | --- |
| $S_1$ | $P_1$ | $(S_1, S_9, S_2, S_{10}, S_3, S_8, S_7, S_5)$ |
| $S_2$ | $P_2$ | $(S_2, S_{10}, S_3, S_8, S_7, S_5)$ |
| $S_3$ | $P_3$ | $(S_3, S_8, S_7, S_5)$ |
| $S_4$ | $P_4$ | $(S_4, S_{11}, S_{10}, S_3, S_8, S_7, S_5)$ |
| $S_5$ | $P_5$ | $(S_5, S_5)$ |
| $S_6$ | $P_6$ | $(S_6, S_1, S_9, S_2, S_{10}, S_3, S_8, S_7, S_5)$ |
| $S_7$ | $P_7$ | $(S_7, S_5)$ |
| $S_8$ | $P_8$ | $(S_8, S_7, S_5)$ |
| $S_9$ | $P_9$ | $(S_9, S_2, S_{10}, S_3, S_8, S_7, S_5)$ |
| $S_{10}$ | $P_{10}$ | $(S_{10}, S_3, S_8, S_7, S_5)$ |
| $S_{11}$ | $P_{11}$ | $(S_{11}, S_{12})$ |
| $S_{12}$ | $P_{12}$ | $(S_{12}, S_{11}, S_{10}, S_3, S_8, S_7, S_5)$ |
| $S_{13}$ | $P_{13}$ | $(S_{13}, S_6, S_1, S_9, S_2, S_{10}, S_3, S_8, S_7, S_5)$ |

The combination of these paths results in the cellular state transition path tree, (*S*_9_, *S*_10_, *S*_8_, *S*_11_, 0, *S*_1_, *S*_5_, *S*_7_, *S*_2_, *S*_3_, *S*_10_, *S*_11_, *S*_6_). We know the state of the winner of each map grid and we use the KNN method to classify the remaining cells based on Euclidean distance to the nearest three winners. Then we smooth the data and produce the cellular state map for all cells (Figure 3E) (See details in Methods).

To study the dynamics of gene expression during the transitions, we calculate the pseudotime ordering of the cells, which is shown along the cellular state transition paths (Figure 4A) (See details in Methods). The colors of the cells represent the expression levels of CDX2. The variances of the expression levels of CDX2 in cellular states *S*_2_, *S*_9_, *S*_1_, *S*_6_, and *S*_13_ are smaller than the ones in other cellular states. The greater variances might be one of the reasons of resulting in the cellular state transition branches, *S*_10_*→S*_2_, *S*_10_*→S*_11_ and *S*_11_*→S*_4_, *S*_11_*→S*_12_ (Figure 4A). The cellular state transition trajectory of CDX2 shows that the expression levels of CDX2 in the trophectoderm (TE) is greater than in the inner cell mass (ICM) which includes the primitive endoderm (PE) and epiblast (EPI), consistent with the previous study (Jedrusik *et al.*, 2008). We define a gene as state-driven when its differential expression between the two adjacent states has a p-value of less than 5% as computed by t-test. Pathway enrichment is conducted using the Enrichr tool (See details in the Supplementary file). Briefly, Fisher’s exact test is used to obtain a p-value for the number of pathway members present among the state-driven genes defined in our experiment. If the pvalue falls within our critical region (5%), we determine the pathway to be significantly enriched. We present the pathway enrichment analysis for the cellular state transitions *S*_10_*→S*_2_ (Figure 4C). They illustrate that PluriNetWork, ESC pluripotency pathway, regulation of actin cytoskeleton pathway and focal adhesion-PI3K-Akt-mTOR-signaling pathway involve significantly in the mouse early embryo development (Zhao and Guan, 2011; Reiske *et al.*, 1999). For the first transition branch *S*_10_*→S*_2_, *S*_10_*→S*_11_, we can see that the transition probability of *S*_10_*→S*_11_ is 19% and the transition probability of *S*_10_*→S*_11_ is 18%. Additionally the replication probability of *S*_2_ is 67% and the replication probability of *S*_11_ is 54%. This provides a clear mechanism for the difference in population size between *S*_2_ and *S*_11_. Interestingly, we find the transition probability of *S*_11_*→S*_4_, 28%, is greater than the one of *S*_11_*→S*_12_, 18%. And the replication probability of *S*_4_ is also greater than the one of *S*_12_. However, the number of the cells in the cellular state *S*_4_ is smaller than the one in the cellular state *S*_12_. Perhaps the replication rate of cells in *S*_12_ is higher, despite a smaller replication probability for each individual cell, due to a much shorter cell cycle (Kelly *et al.*, 1978; Artus and Cohen-Tannoudji, 2008).

**Figure 4.**
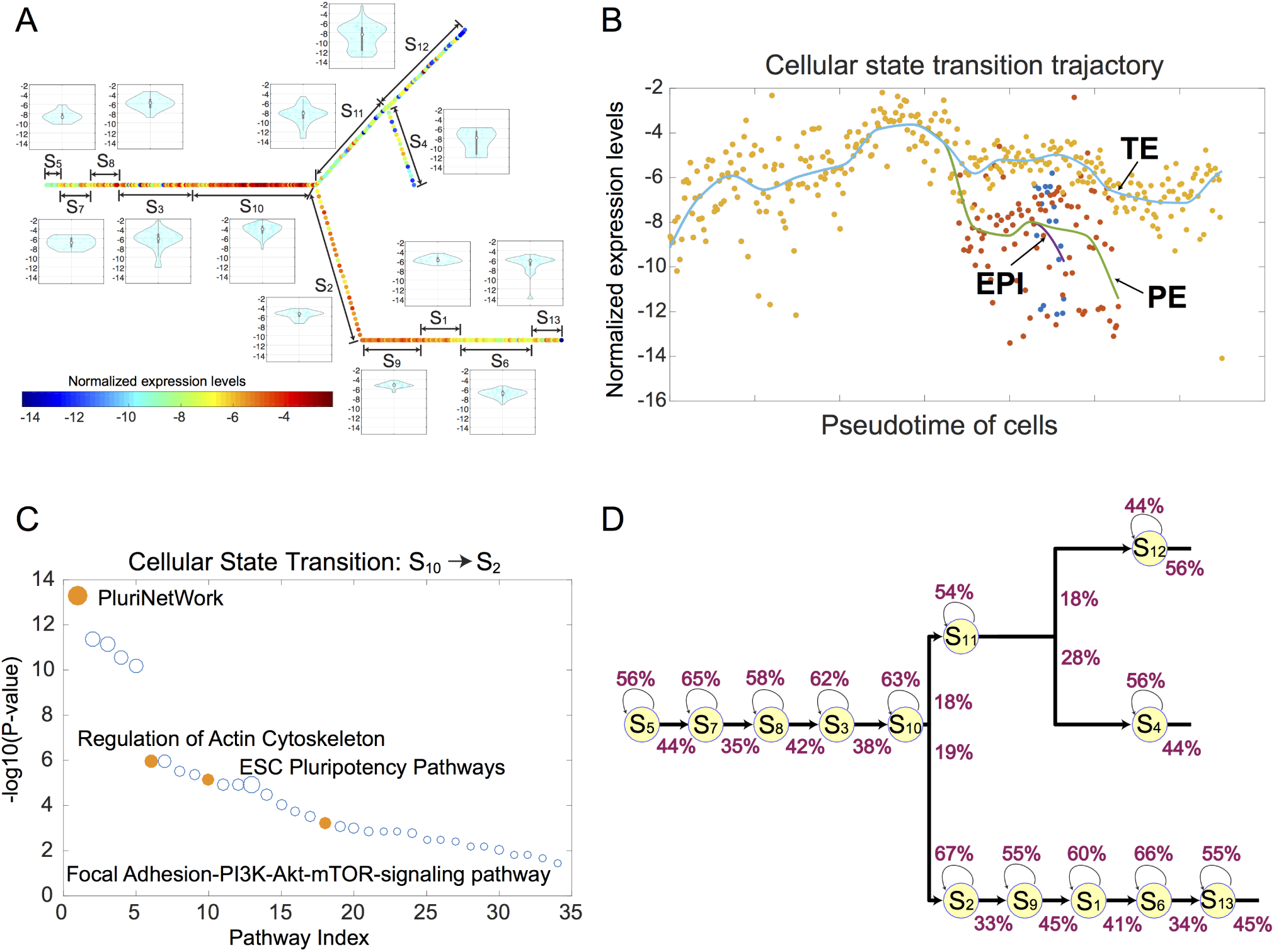
The dynamics of gene expression levels during the cellular state transitions and pathway enrichment analysis of the cellular state transitions. (A). The pseudotime ordering of cells. The colors represent the expression levels of CDX2. The violin plots are the distributions of the expression levels of CDX2 in each cellular state. (B) The cellular state trajectory of CDX2. (C) The bubble charts of pathway enrichment for different cellular state transitions, *S*_10_*→S*_2_. The x-axis is the pathway index and y-axis is –*log*_10_(P-value). Each circle is one pathway. The pathways related with the data are highlighted. (D) The cellular state transition map. The percentage numbers present the probability of the cellular state transition replication and cellular state transitions.

We validate the results calculated by SOMSC against our prior knowledge of the cells temporal order and differentiation trajectory. The temporal ordering of the cellular states identified by SOMSC is consistent with the stage information of the cells in the data (Figure S10). The cells in the cellular state *S*_6_, *S*_7_, *S*_8_, *S*_3_, and *S*_10_ are collected at the 1-cell stage, the 2-cell stage, the 4-cell stage, the 8-cell stage, the 16-cell stage respectively. The cells harvested at the 32-cell stage are located in the areas of the cellular states *S*_2_ and *S*_11_ in the cellular state map and the ones harvested at the 64-cell stage are at the regions of the cellular states *S*_4_, *S*_12_, *S*_9_, *S*_1_, *S*_6_, and *S*_13_ in the map. Finally, SOMSC can track the two state transition branches during the mouse early embryo development (Zernicka-Goetz *et al.*, 2009; Pedersen *et al.*, 1986). The results calculated by SOMSC show that those two fate decisions occur at the 32-cell stage and the 64-cell stage.

### 3.3. SOMSC on scRNA-seq data of mouse haematopoietic stem cell differentiation

To analyze discrete genomic states and the transitional intermediates that span myelopoiesis, the previous study performed single-cell RNA sequencing (scRNA-seq) on 382 cells consisting of stem/multipotent progenitor cells, common myeloid progenitor (CMP) cells, granulocyte monocyte progenitor (GMP) cells, and LKCD34+ cells that includes granulocytic precursors (Olsson *et al.*, 2016). The quality control was performed in the original dataset and therefore the data of those 382 cells are used as the input to SOMSC (Olsson *et al.*, 2016). Out of 23955 genes measured in the original data, 1240 highly variable genes were selected.

We use the data from 382 cells to produces the topographic chart by SOM (Figure S11). We identify 14 basins by the watershed algorithm, labeled *C*_1_, *C*_2_, *…*, *C*_14_ (Figure S11 and S12). The heights of the barriers between adjacent basins are the red numbers on the yellow barriers (Figure S12). The t-test is performed gene-by-gene between the cells of each basin-pair, *C*_*i*_ and *C*_*j*_ and then we calculate the number *T*_*ij*_ of the genes, which are not differentially expressed at 5% significance. Based on all calculated *T*_*ij*_, we merge the following pairs to generate a new topographic chart: (*C*_1_*, C*_5_), (*C*_6_*, C*_7_), (*C*_7_*, C*_9_), (*C*_10_*, C*_11_), (*C*_1_*, C*_13_) (Figure S12). The basins in the topographic chart are labeled *S*_1_, *S*_2_, *…*, and *S*_9_ and each basin represents a cellular state. The distance between the adjacent cellular states in the chart is shown in red (Figure S13 and S14) and used to construct the cellular state network (Figure 5A).

**Figure 5.**
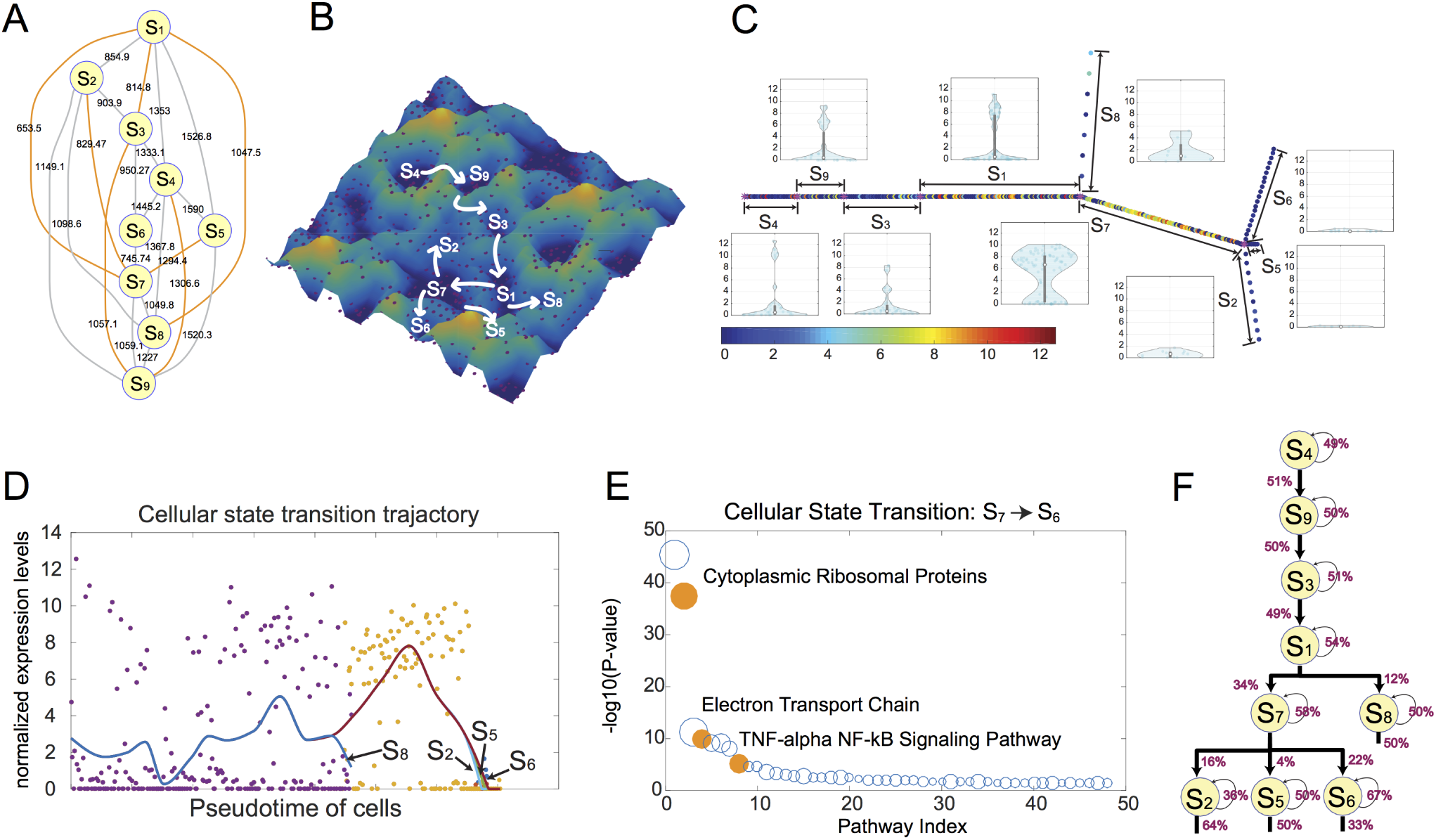
SOMSC reconstructed the cellular state transition path using the scRNA-seq data of mouse haematopoietic stem cell differentiation (Moignard *et al.*, 2013). (A) The cellular state network is built based on the topographic chart. The nodes are the cellular states. The weight of the edge is the height of the barrier of adjacent basins. The orange line is the edge with the smallest weight associated with each cellular state. (B) The cellular state map. The red balls are the cells. The basins correspond to the cellular states. (C) The pseudotime ordering of cells. The colors represent the expression levels of Irf8. The violin plots are the distributions of the expression levels of Irf8 in each cellular state. (D) the cellular state trajectory of Irf8. (E) the bubble charts of pathway enrichment for different cellular state transitions, *S*_7_*→S*_6_. The x-axis is the pathway index and y-axis is –*log*_10_(P-value). Each circle is one pathway. The pathways related with the data are highlighted. (F) The cellular state transition path tree with the cellular state replication probability and the cellular state transition probabilities.

The combination of these paths *P*_1_, *P*_2_, *…*, and *P*_9_ results in the cellular state transition path tree, (*S*_3_, *S*_7_, *S*_9_, 0, *S*_7_, *S*_7_, *S*_1_, *S*_1_, *S*_4_). The cellular state map including all 382 cells was generated as previously described (Figure 5B). SOMSC identifies nine cellular states, which is consistent with previously reported classification of myelopoetic cells based on scRNA-seq (Olsson *et al.*, 2016). Importantly, SOMSC faithfully recovers the differentiation trajectories of the cells: cellular states *S*_4_, *S*_9_, *S*_3_ are the LSK cells and *S*_1_ consists of CMP and GMP cells. There is a branch from the cellular state *S*_1_ to the cellular state *S*_7_ and to the cellular state *S*_8_. All cells in the cellular state *S*_8_ are CMP cells and the ones in the cellular state *S*_7_ are GMP cells. We find that Irf8 is expressed in the cellular state *S*_7_ (Figure 5C). The cells in the cellular state *S*_2_ are CMP, the ones in the cellular state *S*_5_ are LSKs and the ones in the cellular state *S*_6_ are GMP cells. From the violin plots in Figure 5C we can see that there is more than one mode in the cellular states *S*_4_, *S*_9_, *S*_3_, *S*_1_, and *S*_7_, suggesting that the cell fate decision is made relatively early.

In order to determine the genes driving the cellular state transitions we identify state-driven genes by t-test as previously described. Based on the list of differentially expressed genes, the enriched pathways are obtained by gene set enrichment as previously described. We present the pathway enrichment analysis for the cellular state transitions *S*_7_*→S*_6_ (Figure 5E), indicating that TNF*α*-NF*?*B pathway is critical during the differentiation of GMP cells (See more results in the Supplementary file).

Next we estimate the replication probabilities and transition probabilities of the cellular states (Figure 5F). There are two branches in this cellular state transition path tree. The results show that the transition probability from *S*_1_ to *S*_7_ is 34% and the transition probability from *S*_1_ to *S*_8_ is 12%. The latter probability is smaller, corresponding to fewer cells classified in *S*_8_ than in *S*_7_. The results also show that the probability of the transition from *S*_7_ to *S*_6_, *S*_7_ to *S*_2_, and *S*_7_ to *S*_5_ are 22%, 16%, and 4% respectively. Correspondingly, *S*_6_ represents the largest population among all three fates.

The cellular state transition path tree calculated by the SOMSC is consistent with the hematopoietic hierarchy. *S*_4_ is the long-term HSC. *S*_9_ is the short-term HSC. *S*_3_ is the multipotent progenitor. *S*_1_ is the common myeloid progenitor. *S*_7_ is the granulocyte-macrophage progenitor. *S*_8_ is the megakaryocyte-erythrocyte progenitor (Figure S15). Since Gfi1 is expressed only in the cells of *S*_6_ among *S*_6_, *S*_5_, and *S*_2_ (Figure S16), then *S*_6_ is the granulocyte cell. Because Irf8 is highly expressed in the cells of *S*_2_ (Figure 5C), then the cells in *S*_2_ are the monocyte cells. We predict that the cells in *S*_5_ are the dendritic cells. All of them need to be verified.

### 3.4. SOMSC on scRNA-seq data of adult mouse olfactory stem cell lineage trajectories

616 cells were collected to define a detailed map of the postnatal olfactory epithelium showing the cell fate potentials and branchpoints in olfactory stem cell lineage trajectories by whole transcriptome profiling scRNA-seq (Fletcher *et al.*, 2017). We select 824 highly variable genes out of 42127.

As before, we use an SOM to produce a topographic chart using the expression profiles of all 616 cells (Figure S17). We identify 22 basins by the watershed algorithm, labeled *C*_1_, *C*_2_, *…*, *C*_14_ (Figure S18). Then 17 cellular states are determined by SOMSC (see Methods) and their corresponding cellular state network is constructed (Figure S19 and S20). Next, the CSM and the cellular state transition path are obtained (Figure 6). To validate our results, we utilize the prior categorization of these cells, which are available from (GEO accession GSE95601) (Fletcher *et al.*, 2017). They include the resting HBCs, immediate Neuronal Precursor 1 (INP1), Globose Basal Cells (GBCs), mature Sustentacular Cells (mSUS), transitional HBC 2, immature Sustentacular Cells (iSUS), transitional HBC 1, immature Olfacotry Sensory Neurons (iOSNs), Immediate Neuronal Precursor 3 (INP3), Microvillous Cells, type 1, mature Olfactory Sensory Neurons (mOSNs), Immediate Neuronal Precursor 2 (INP2), and Microvillous Cells (MV) (Fletcher *et al.*, 2017). The cellular states are denoted by the cluster labels used in the original data. The non-differentiating cell types are determined by the known marker genes (mature olfactory sensory neurons, mature sustentacular cells, and microvillus cells) (Fletcher *et al.*, 2017). All of these cellular types correspond to the cellular states, which are the endpoints in the cellular state transition paths, consistent with our prior knowledge (Figure S21). Notably, SOMSC identifies the sustentacular cluster as the endpoint in the cellular state transition path tree without using the prior knowledge (Figure 6CF) not possible using previous methods (Fletcher *et al.*, 2017). The SOMSC can not only identify those three primary cellular state transition paths but also detect more cellular state transition paths than what previous methods obtained (Figure 6CF). Interestingly, other lineage detection methods fail to recognize the three primary lineages in this dataset (Street *et al.*, 2017). We suggest here that GBC is categorized as an intermediate state, and a precursor cellular state of the immediate neuronal precursors. And that immature sustentacular cells may be the progenitor cells for mature olfactory sensory neurons. In addition, transitional horizontal basal cells are found in different branches. Finally, the microvillus cells are recognized as a separate branch different from the neuronal lineage, which is consistent with the previous results (Fletcher *et al.*, 2017).

**Figure 6.**
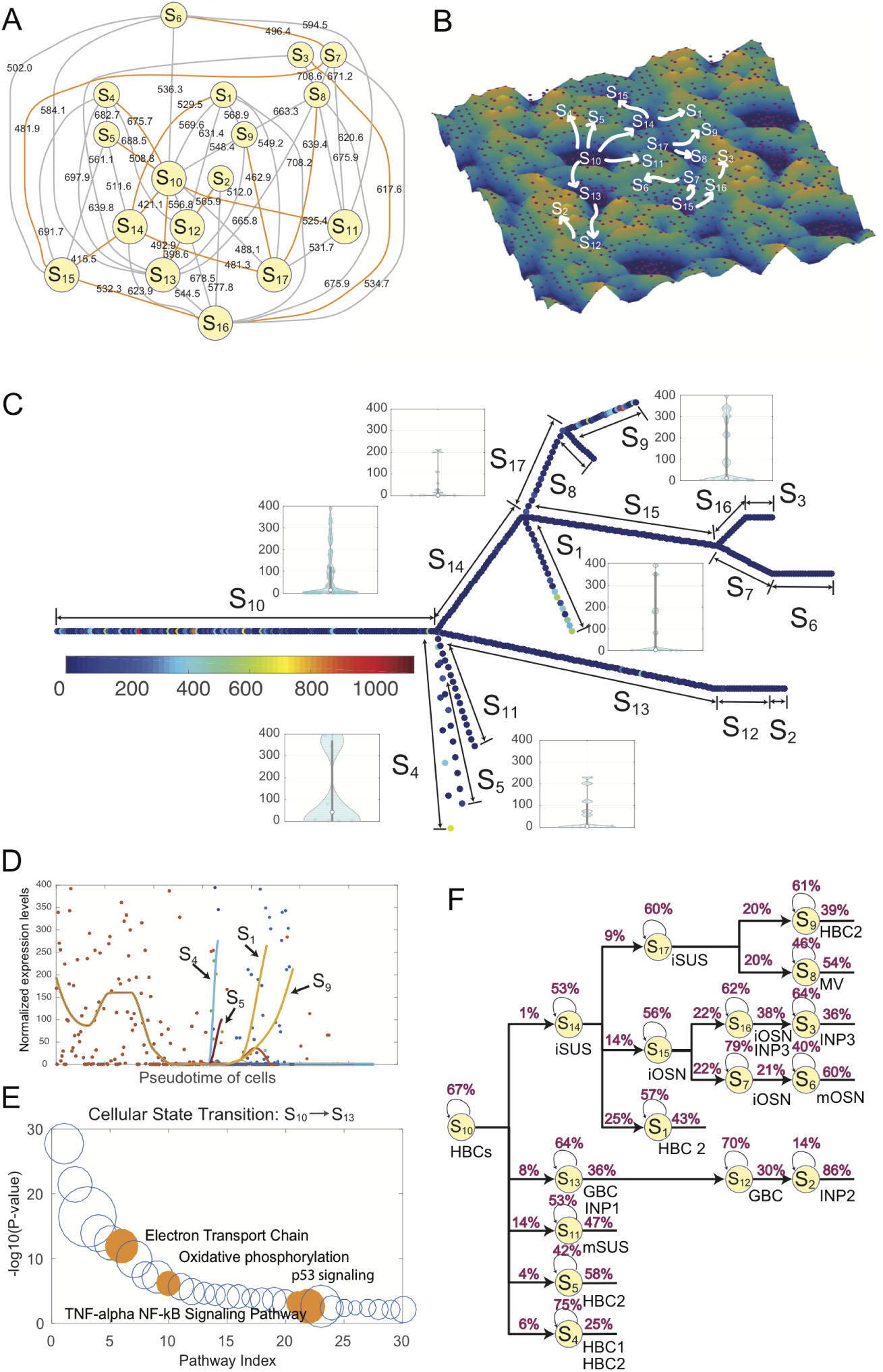
SOMSC reconstructed the cellular state transition path using the scRNA-seq data of adult mouse olfactory stem cell lineage trajectories (Fletcher *et al.*, 2017). (A) The cellular state network is built based on the topographic chart. The weight of the edge is the height of the barrier of adjacent basins. The orange line is the edge with the smallest weight associated with each cellular state. (B) The cellular state map. The red balls are the cells. The basins correspond to the cellular states. (C) The pseudotime ordering of cells. The colors are quantified by the expression levels of Trp63. The violin plots are the distributions of the expression levels of Trp63 in each cellular state. (D) the cellular state trajectory of Trp63. (E) the bubble charts of pathway enrichment for different cellular state transitions, *S*_10_*→S*_13_. The x-axis is the pathway index and y-axis is –*log*_10_(P-value). Each circle is one pathway. The pathways related with the data are highlighted. (F) The cellular state transition path tree with the cellular state replication probability and the cellular state transition probabilities.

An interesting observation from the cellular trajectory of gene Trp63 shows that the expression levels of cells in *S*_10_ (HBC1) change from high to low and they become rather low before the cells transit into other cellular states, which was also observed in the previous study (Schnittke *et al.*, 2015) (Figure 6D). The pathway enrichment analysis indicates that p53 signaling pathway plays an important role during the cellular transition from HBCs to GBCs, which is consistent with the previous observations (Herrick *et al.*, 2017) (Figure 6E). It also shows that the pseudotime ordering of cells generated by SOMSC is consistent with previous study (Schnittke *et al.*, 2015). The estimated values of replication probability and the transition probabilities suggest that the length of cell cycle may play a critical roles at some cellular transitions. For example, the transition probability from *S*_10_ to *S*_4_ (6%) is very close with the one from *S*_10_ to *S*_13_ (8%). The value of replication probability of *S*_13_ is smaller than the one of *S*_4_ (Figure 6F). However, the number of cells in the cellular state *S*_13_ is much greater than the one in the cellular state *S*_4_ indicating that the shorter cell cycle length of cells in the cellular states may be one of the key factors during the development of olfactory (Huard and Schwob, 1995).

## 4. Conclusion and Discussion

In this work we have presented a SOM based method for analyzing gene expression data of single cells that may contain multiple cellular states. Applications of SOMSC to a set of simulated data and three sets of experimental data have demonstrated the capabilities and effectiveness of SOMSC in identifying cellular states and their cellular state transition path trees.

The CSM based on the cellular states identified by SOM provides a global landscape view of the cellular states and their transitions. The location of each cell in the CSM may provide useful information on the cells viability, potential of transitions to different cellular states and the replication/transition probabilities using the unbiased selection of genes. These properties make our algorithm unique compared with many other methods for single-cell analysis.

The major computational cost of SOMSC comes from iteratively computing the U-matrix in the SOM, which has a complexity of 𝒪(*N N*_*g*_*D T*) where *D* is the number of genes measured in the data, *T* is the number of iterations used in SOM, and *N* is the number of cells in the data set(Lee and Verleysen, 2007). In practice, *D* is usually less than 10,000 (the number of genes significantly expressed), and both *T* and *N* are less than 1,000, implying an average complexity of 𝒪(10^10^).

The one parameter of SOMSC is the map size. A map with too many grids may produce too many small clusters while too few grids may lead to a map containing too few clusters (Figure S14A, S14B, S14C). In our analysis, we set the number of grids approximately equal to the number of cells in each dataset.

Single-cell data is often used to identify cellular states in heterogeneous cell populations (Kolodziejczyk *et al.*, 2015). SOMSC can capture the shape of data which lacks the convex or normal structure required by many other methods (Liebscher *et al.*, 2012). Another major feature of SOM is the identification of multiple minima since SOM searches the entire space of feasible solutions during early exploration, and divides the search space gradually until it finds an optimal solution (Liebscher *et al.*, 2012; Openshaw *et al.*, 1995). When SOM and k-means use the same initial guess near the optimal solution to approach the optimal, similar performance and similar results are obtained(Liebscher *et al.*, 2012; Openshaw *et al.*, 1995). This is consistent with the observation that SOMSC is rather stable in finding the basins of attraction and the transition paths from the CSM of the single-cell data.

Previous work has shown that confounding factors (e.g. batch errors, cell cycle effects) impede analysis of single-cell data (Buettner and Theis, 2012; Buettner *et al.*, 2015). PCA (Pickrell *et al.*, 2010), surrogate variable analyses (Leek and Storey, 2007), probabilistic estimation of expression residuals (Stegle *et al.*, 2010, 2012) and factor analysis (Risso *et al.*, 2014) have been explored to reduce the effects of confounders in gene expression studies on bulk cell populations (Stegle *et al.*, 2015). Most of these methods could be extended to the data of single cells, however, removal of cell cycle effects, an important source of variation in single-cell measurement, is still challenging. However, a linear mixed model has recently been utilized to remove cell-cycle dependent gene expression as a source of variation (Buettner *et al.*, 2015).

The CSM in many ways is similar to the landscape description although the typical landscape is a function of each gene. It would be interesting to make a comparison between a landscape computed by forward modeling based on a small size of network and the CSM generated by data. In general, SOMSC is a robust method, which is also convenient for visualization, to identify the cell states based on single-cell data. It facilitates the identification of pathway components, such as signature transcription factors, and pseudo temporal ordering of cells involving complex differentiation trajectories.

## Acknowledgement

We are grateful to Matt Karikomi for manuscript proofreading.

